# Machine learning assistive rapid, label-free molecular phenotyping of blood with two-dimensional NMR correlational spectroscopy

**DOI:** 10.1101/2020.06.20.162974

**Authors:** Weng Kung Peng, Tian-Tsong Ng, Tze Ping Loh

**Author notes:** **Corresponding authors:** W. K. Peng, T. P. Loh.

## Abstract

Translation of the findings in basic science and clinical research into routine practice is hampered by large variations in human phenotype. Developments in genotyping and phenotyping, such as proteomics and lipidomics, are beginning to address these limitations. In this work, we developed a new methodology for rapid, label-free molecular phenotyping of biological fluids (e.g., blood) by exploiting the recent advances in fast and highly efficient multidimensional inverse Laplace decomposition technique. We demonstrated that using two-dimensional T_1_-T_2_ correlational spectroscopy on a single drop of blood (<5 μL), highly time– and patient–specific ‘molecular fingerprint’ can be obtained in minutes. Machine learning techniques were introduced to transform the NMR correlational map into user-friendly information for point-of-care disease diagnostic. The clinical utilities of this technique were demonstrated through the direct analysis of human whole blood in various physiological (e.g., oxygenated/deoxygenated states) and pathological (e.g., blood oxidation, hemoglobinopathies) conditions.

## Introduction

High-resolution nuclear magnetic resonance (NMR) spectroscopy is a powerful and attractive technique in biochemistry (e.g., for structural protein analysis^1^, characterizing metabolomics responses in biological samples^2–4^) and inorganic chemistry^5^. However, high-resolution NMR systems are large, expensive and incompatible with in-situ or portable applications. There is an increasing demand for low field portable NMR system for use in food sciences^6^, oil-gas exploration^7^, and point of care clinical testing^8–10^. In high field NMR, biochemical information is typically detected and encoded in the frequency domain (“chemical shift”), in which the spectral resolution scale with respect to the external magnetic field. This reduces its portability and limit its downstream application in a large scale manner.

However, biochemical and biophysical information (e.g., molecular rotational, diffusional motion) can also be encoded in the relaxation times frame, namely the longitudinal (T_1_) and transverse (T_2_) using a portable low field NMR system. In addition, molecular information in the time-domain can be inversely decoded with the availability of fast and reliable Laplace inversion algorithm^7,11^. This can provide parallel information that is not available in the traditional NMR frequency domain based spectra.

In recent years, significant advances in NMR system miniaturization^8,12–14^ (e.g., electronic console^12,13,15,16^, radio-frequency probe^9,10,17–19^, microfluidic-based chip^20,21^) utilizing small foot-print permanent magnetic (<1 Tesla) for one-dimensional NMR relaxometry on water-proton (e.g., spin-spin relaxation (T_2_-relaxation)) have been widely applied for point-of-care medical testing^8,9,13^. These include immuno-magnetic labelled (e.g., tumour cells^8,21^, tuberculosis^22^ and magneto-DNA detection of bacteria^23^) and the label-free detection of various pathological states such as oxygenation^19^/oxidation level^10^ of the blood, malaria screening^9,24^, and rapid phenotyping of oxidative stress in diabetes mellitus^25,26^ (Supplementary Table 1).

We demonstrated the first unique two-dimensional ‘molecular fingerprint’ of a single drop of blood (<5 μL) obtained in minutes (Supplementary Fig. 1) using two dimensional T_1_-T_2_ correlational spectroscopy with an inexpensive, bench top sized NMR spectrometer (Fig. 1). By exploiting the recent development of fast and highly efficient multidimensional inverse Laplace decomposition algorithm^7,27^, unique two-dimensional signature of various haemoglobin (Hb) derivatives with respect to its magnetic resonance relaxation reservoirs in oxygenated (oxy-Hb), deoxygenated (deoxy-Hb) and oxidized (oxidized Hb) states were observed for the first time and its phenotypic expression in various pathological states (e.g., blood oxidation, hemoglobinopathies) are reported in this work. Machine-learning techniques (e.g., multidimensional scaling (MDS), t-SNE, Isomap) were introduced to transform the NMR correlational maps into user-friendly information for medical decision making (Fig. 2). We report that the supervised models (e.g., neural network) were at least on par or outperformed the average trained human being in performing deep image analysis (Table 3).

**FIG. 1:**
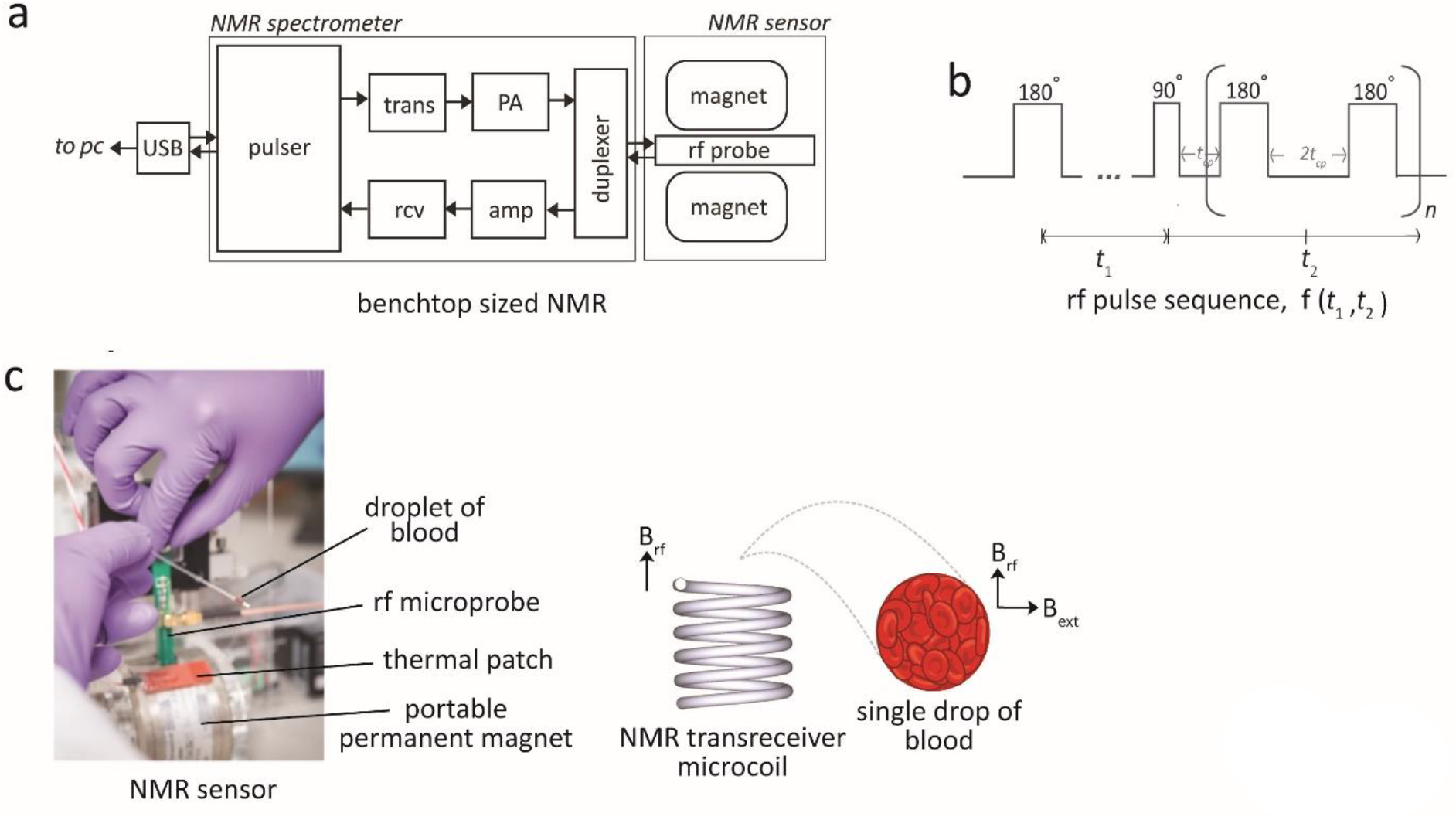
Two-dimensional NMR T_1_-T_2_ correlational spectroscopy for molecular phenotyping of blood. (a) Schematic diagram of the bench-top sized NMR-based POCT system. The applied radio frequencies were centred at 21.57 MHz, which corresponds to the Larmor frequency of water-proton in 0.5 Tesla of the permanent magnet. The 90-degree pulse used is 10 μs. The whole system is lightweight (<2kg) and portable suitable for in-situ measurements. The abbreviations are; USB: Universal Serial Bus, trans: Transmitter, rcv: Receiver, amp: pre-amplifier, PA: power amplifier, rf: radio-frequency, and PC: personal computer. (b) The pulse sequence used for the T_1_-T_2_ correlational spectroscopy is the modified inversion recovery with CPMG observation. It is encoded for a period of *t*_1_ and subsequently spaced for a period of *t*_2_ for n-train pulses, in entirely in analogous to the two-dimensional NMR spectroscopy in the frequency domain. The relaxation properties can be used as a highly sensitive and specific molecular probe, and provide important molecular motion (e.g., correlational relaxation, diffusion properties), which is not readily available in NMR spectra in the frequency domain. (c) A single drop of whole blood contained in a micro capillary tube was spun using standard haematocrit centrifuge (6000*g*, 1 min) to separate and concentrate the RBCs from the plasma. The capillary tube is then loaded into a permanent magnet. The radio frequency coil (inner diameter of 1.20 mm) was adjusted to focus on the packed RBCs.

**Fig. 2:**
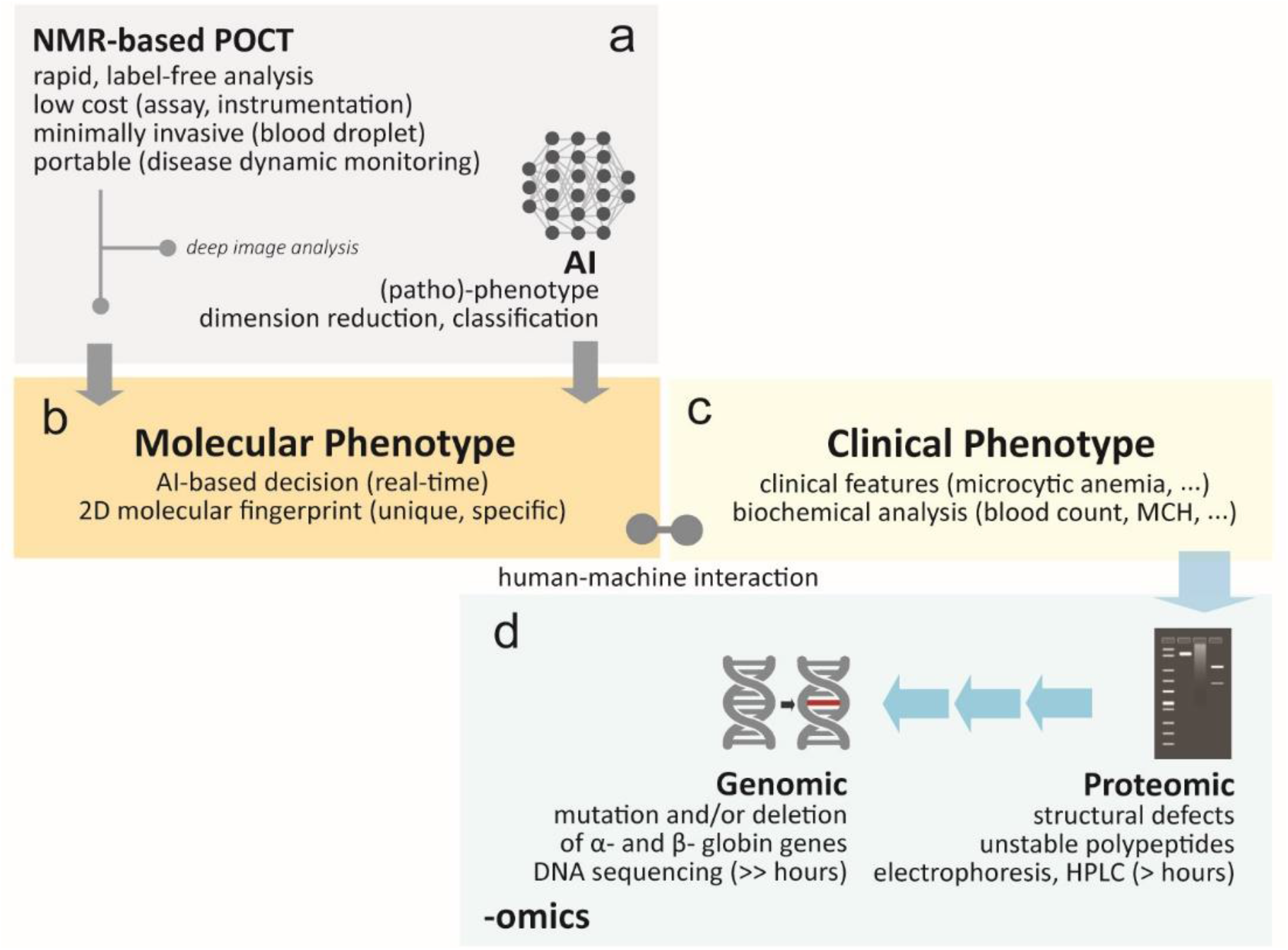
A proposed scheme of human-machine interaction for rapid, label-free disease detection in clinical hemoglobinopathies. (a) The NMR-based POCT is used with (or without) the assistant of artificial intelligence (AI). (b) The highly unique and detailed 2D magnetic resonance based molecular fingerprint can be used directly (without AI) for rapid screening. (c) Clinical phenotype (e.g., clinical representation) can be bias due to subjective human judgment. With AI, deep image analysis (e.g., hierarchical clustering, dimension reduction) were performed to transform the highly complicated data (e.g., hyper dimension) into human friendly information to assist in medical decision making (e.g., diagnostic, staging) in real-time mode (Fig. 6 and Fig. 7). (d) Multi-omics information (e.g., proteomics, genomics) may be performed simultaneously to confirm the genetic variants and/or other anomalies. Back-end laboratory and time consuming test (e.g., high performance liquid chromatography (HPLC)) may be by-passed depending on the outcome of the molecular phenotyping.

## Results

### Water-protein interactions in blood microenvironment

Freshly collected whole blood samples containing predominantly the oxy-Hb were collected from healthy donors (‘wild-type’). Oxygenation and re-oxygenation was achieved with rigorous pipetting in ambient air. Using microcapillary tube, the whole blood was sampled and spun (6000*g*, 1 min) into narrowband of red blood cells (RBCs) for micro NMR measurements.

Three peaks (R-peak, S-peak and T-peak) with (T_2_=141ms, T_1_=562ms), (T_2_=4.47ms, T_1_=335ms) and (T_2_=1.12ms, T_1_=188ms) respectively were observed from the T_1_-T_2_ correlational spectroscopy performed on the water-proton nuclei (^1^H) of the RBCs (Fig. 3a). It appeared that RBCs microenvironment could be decomposed into two major relaxation reservoirs, consisting of one slow relaxation component (R-peak), and two fast relaxation components (S-peak, T-peak), attributed to the interaction of the water molecules with its’ respective microenvironment i.e., bulk water, intermediate hydration layer, macromolecules protein, respectively. Water molecules are subjected to diverse dynamic processes as a result of their interaction with variety of sites/functional groups.

**FIG. 3:**
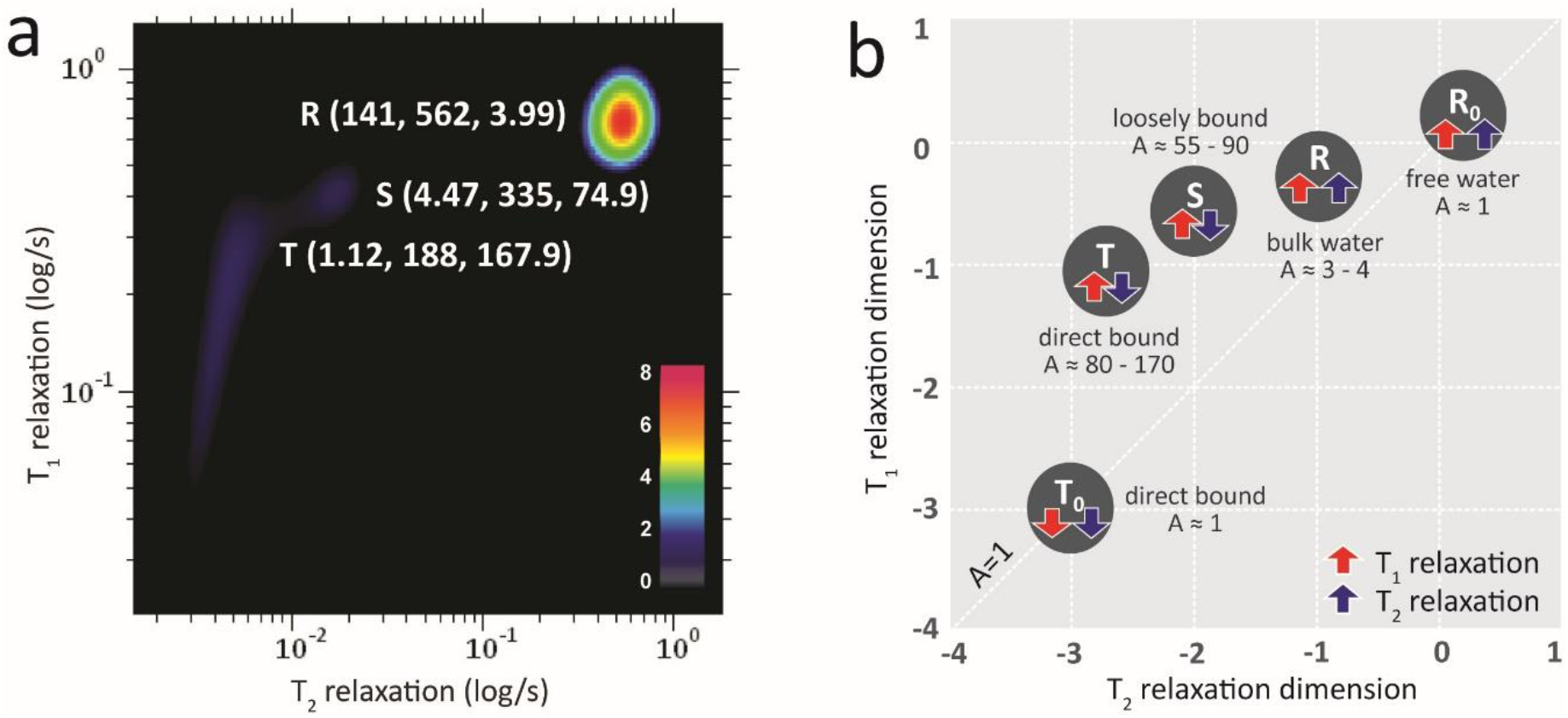
The T_1_-T_2_ correlational spectrum of blood microenvironment. (a) The decomposed relaxation reservoirs (R-peak, S-peak, and T-peak) of packed red blood cells microenvironment with the haemoglobin in oxygenated state. The coordinate is represented as (T_2_ relaxation (in ms), T_1_ relaxation (in ms), A-ratio (unitless)). A-ratio is the ratio between T_1_/T_2_. (b) The multiple relaxation reservoirs of the blood microenvironment in the T_1_-T_2_ correlational spectrum in log-log plot; i.e., the bulk water (R-peak), hydration layer (S-peak), and direct macromolecular protein interaction (T-peak) for haemoglobin in oxygenated state. In the oxidized state, the T-peak dropped substantially (T_0_-peak). The unbound molecule, R_o_ (e.g., free water) located on the diagonal line (A-ratio approaches unity).

The significantly large signal intensity (and slowest relaxation component) of R-peak is attributable to bulk water molecules which makes up more than 98% of the total mass-ratio of RBCs. The bulk water has minimal and indirect contact with macromolecules protein (through long range dipolar couplings), and hence the weakest water-protein interactions. The relaxation dephasing system came predominantly from the dipole-dipole homonuclei coupling of water-to-water network. On the other hand, the presence of two distinct individual peaks (i.e., S-peak, T-peak) suggested that the fast relaxation component can be further resolved into sub-regions^28,29^. The S-peak is the water molecules at the intermediate hydration layer, and the T-peak are water molecules, which came into direct contact with the surface of macromolecular protein. Dortch RD *et al.* and McDonald PJ *et al.*, proposed the idea of exchange peaks^30^ and surface relaxation^31^, respectively, but the observation in this work is in consistent with the three peaks model proposed by Thompson B.C. et al. and Lores G. et al.^32,33^.

Interestingly, each peak (R-peak, S-peak, T-peak) possess consistent and yet unique ratio of T_1_/T_2_ of (3.99, 74.90, 167.86), respectively, which appeared to characterize the degree of water-protein interactions (Fig. 3b and Table 1). We define here the T_1_/T_2_ ratio as A-ratio. With increased water-protein interactions, the motion of water-proton was drastically slower and restricted (and hence the reduced T_1_ relaxation and T_2_ relaxation). The spin-spin relaxation appeared to be much more efficient (shorter T_2_ relaxation) relative to its’ spin-lattice relaxation counterpart and hence a large A-ratio. In contrast, an unbound free molecules in the extreme fast motion region, possess large T_1_ relaxation and T_2_ relaxation, with A-ratio approaches unity (~1). Importantly, the relaxation profile forms unique and specific two-dimensional ‘molecular fingerprint’ of each individual (others wild-type in Supplementary Fig. 3) that is very sensitive to its’ molecular microenvironment measureable at the NMR relaxation timescales.

**Table 1:** The decomposed relaxation reservoirs of packed red blood cells with the haemoglobin in (a) oxygenated, (b) oxidized, and (c) deoxygenated states. A-ratio is the ratio between T_1_/T_2_.

| bulk water, R-peak | T<sub>1</sub> (ms) | T<sub>2</sub> (ms) | A-ratio |
| --- | --- | --- | --- |
| a. oxygenated state | 562 | 141 | 3.99 |
| b. oxidized state | 217 | 120 | 1.81 |
| c. deoxygenated state | 463 | 102 | 4.53 |

| hydration layer, S-peak | T<sub>1</sub> (ms) | T<sub>2</sub> (ms) | A-ratio |
| --- | --- | --- | --- |
| a. oxygenated state | 335 | 4.47 | 74.90 |
| b. oxidized state | 120 | 4.18 | 28.71 |
| c. deoxygenated state | 242 | 2.71 | 89.30 |

Table 1: The decomposed relaxation reservoirs of packed red blood cells with the haemoglobin in 5 (a) oxygenated, (b) oxidized, and (c) deoxygenated states. A-ratio is the ratio between T<sub>1</sub>/T<sub>2</sub>.
| direct bound macromolecules, T-peak | T<sub>1</sub> (ms) | T<sub>2</sub> (ms) | A-ratio |
| --- | --- | --- | --- |
| a. oxygenated state | 188 | 1.12 | 167.86 |
| b. (i) oxidized state | 50.3 | 1.34 | 37.54 |
| b. (ii) oxidized state | 2.43 | 0.78 | 3.12 |
| c. deoxygenated state | 175 | 0.565 | 309.73 |

### Oxidative degradation of haemoglobin in blood

Freshly collected whole blood sample which consists of predominantly the oxy-Hb was oxidized to oxidized Hb in the presence of sodium nitrite, and spun down for NMR measurements (see Methods Online). The relaxation times of the three major peaks were (R-peak: T_2_=120ms, T_1_=217ms), (S-peak: T_2_=4.18ms, T_1_=120ms), and (T-peak: T_2_=1.34ms, T_1_=50.3ms) in oxidized state reduced considerably as compared to the baseline oxygenated state (non-oxidized, diamagnetic state) (Figs. 4a-b). The presence of excessive oxidized Hb in blood causes serious tissue hypoxia, a pathological state known clinically as methemoglobinemia^34^.

**FIG. 4:**
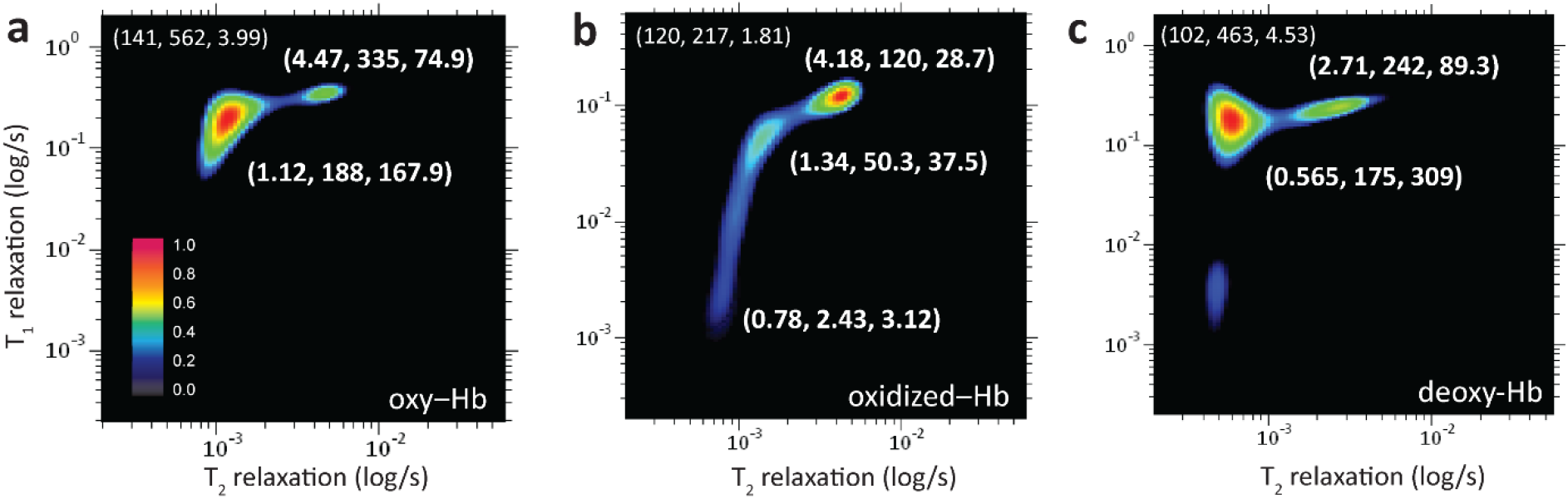
The T_1_-T_2_ correlational spectrum of blood microenvironment of packed red blood cells in (a) oxygenated, (b) oxidized, and (c) deoxygenated states. The zoom-in details of decomposed relaxation reservoirs for fast relaxation components (S-peak and T-peak), while the slow relaxation component (bulk water molecules, R-peak) is not shown. The coordinate for R-peak is indicated at upper left of the spectrum. The coordinate is represented by (T_2_ relaxation (in ms), T_1_ relaxation (in ms), A-ratio). Freshly prepared oxy-Hb was subjected to oxidation with 10 mM sodium nitrite for 45 minutes, and sodium dithionite (in excess) for 40 min to chemically locked the Hb in the deoxygenated state. All the samples were washed thrice and resuspended into 1x PBS for micro MR measurements. The experimental parameters used were echo time=200 μs, T_1_-incremental steps=32 steps, and signal averaging=4. The number of echoes used were 4000 (oxygenated Hb) and 2000 (oxidized Hb deoxygenated Hb).

The marked relaxation enhancement observed was due to the presence of five unpaired electrons in the ferric iron (Fe^3+^), which acted as the paramagnetic relaxation center^34^. The magnetic moment of ferric iron is 1000-fold higher than that of one single proton^34,35^. Significantly, due to the long range dipolar nuclei-electron, the paramagnetism of the unpaired electrons had considerable effect on the bulk water molecules (R-peak). In contrast to the oxygenated states (in diamagnetic state), the spin-lattice relaxation effect in oxidized states (in paramagnetic state) appeared to be much more efficient in comparison to the spin-spin relaxation effect and hence the reduction in A-ratio=1.81 (Table 1).

A distinctively long stretch of T_1_-relaxation distribution, extending across two orders of magnitude (ca., 1ms to 100ms along the T_1_ dimension) displayed by the protein-bound water-proton molecules (from T-peak to T_0_-peak). The ‘relaxation tail’ originating from (T_2_=1.34ms, T_1_=50.3ms) to (T_2_=0.78ms, T_1_=2.43ms), notably became a distinctive feature of oxidized Hb. This is due to the distance (r)-dependent paramagnetism effect, in which the relaxation efficiency reduced at the rate of 1/r^6^ from its relaxation center^34^. As the proton nuclei approach the relaxation centre (of the unpaired electron), the T_1_- and T_2_-relaxation components reduced to a comparable rate (A-ratio approaching unity, T_o_ in Fig. 3b). The gradual process of Hb oxidation under the exposure of mild oxidant were captured in a well-controlled manner confirmed the existence of transitional states in the formation of ‘relaxation tail’ (Supplementary Fig. 2).

On the other hand, the protein-bound water molecules (T-peak) in the deoxygenated states, exhibited profound T_2_ shortening (0.565ms) with relatively very little T_1_ shortening (175ms) due to the short relaxation time of electron and its obscure protein configuration^36^. As a result, the A-ratio of deoxy-Hb (309.7) is distinctively larger than its oxy-Hb (167.9) and oxidized Hb (37.5) counterpart (Fig. 4c).

### Rapid molecular phenotyping in clinical hemoglobinopathies

We demonstrated the clinical utility of molecular phenotyping in clinical hemoglobinopathies by mapping out the spectrum of heterozygous HbE, HbD and a heterozygous beta thalassemia (HBB:c.27_28insG) variants (Fig. 5 and Table 2). An additional six other Hb variants (in Supplementary Fig. 3b) were received for machine learning and blind test studies (Table 3 and Fig. 7). A limitation of this study was that the current study only involve heterozygous HbE phenotype. Given the low prevalence of homozygous HbE variant phenotype (~0.1%) in our population^**37**^, therefore, we were unable to include such subject during the study period. The Hb variants were first identified by a cation-exchange high performance liquid chromatography method (Bio-Rad Variant II analyser) and further confirmed by capillary electrophoresis (Sebia CAPILLARYS 2 analyser) and genotyping. NMR measurements were carried out in its native state (without any chemical treatment) of the spun down packed RBCs.

**FIG. 5:**
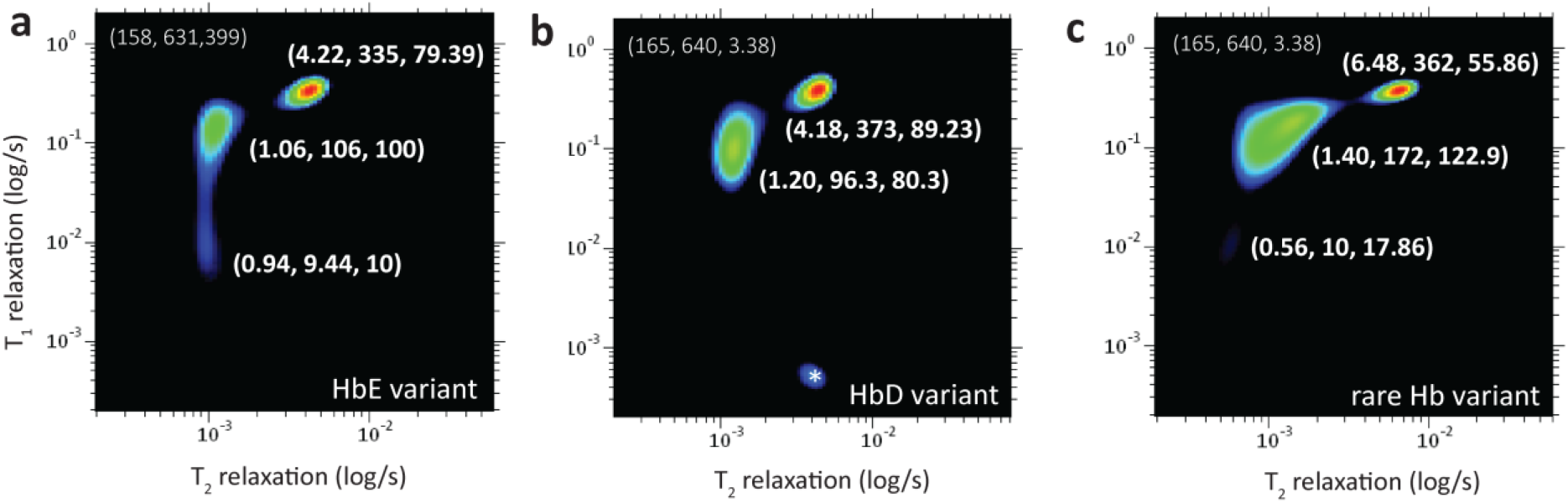
The T_1_-T_2_ correlational spectrum of blood microenvironment in haemoglobin variants in (a) HbE variant, (b) HbD variant, and (c) beta thalassemia variant, and further six other Hb variants (in Supplementary Fig. 3b). The zoom-in details of decomposed relaxation reservoirs for fast relaxation components (S-peak and T-peak), while the slow relaxation component (bulk water molecules, R-peak) is not shown. The coordinate for R-peak is indicated at upper left of the spectrum. The coordinate is represented by (T_2_ relaxation (in ms), T_1_ relaxation (in ms), A-ratio). The experimental parameters used were echo time=200 μs, number of echoes=4000, T_1_-incremental steps=32 steps, and signal averaging=4. Note that there is a possible artefact denoted as (*).

**Table 2:** The decomposed relaxation reservoirs of packed red blood cells with the haemoglobin variants in (a) wild-type (control), (b) HbE variant, (c) HbD variant, and (d) rare beta thalassemia variant.

| bulk water, R-peak | T<sub>1</sub> (ms) | T<sub>2</sub> (ms) | A ratio |
| --- | --- | --- | --- |
| a. Hb wild-type (control) | 562 | 141 | 3.99 |
| b. HbE variant | 631 | 158 | 3.99 |
| c. HbD variant | 640 | 165 | 3.38 |
| d. rare Hb variant | 640 | 165 | 3.87 |

| hydration layer, S-peak | T<sub>1</sub> (ms) | T<sub>2</sub> (ms) | A ratio |
| --- | --- | --- | --- |
| a. Hb wild-type (control) | 335 | 4.47 | 74.94 |
| b. HbE variant | 335 | 4.22 | 79.39 |
| c. HbD variant | 373 | 4.18 | 89.23 |
| d. rare Hb variant | 362 | 6.48 | 55.86 |

| direct bound macromolecules, T-peak | T<sub>1</sub> (ms) | T<sub>2</sub> (ms) | A ratio |
| --- | --- | --- | --- |
| a. Hb wild-type (control) | 188 | 1.12 | 167.9 |
| b. HbE variant | 106 | 1.06 | 100 |
| c. HbD variant | 96.3 | 1.20 | 80 |
| d. rare Hb variant | 172 | 1.40 | 122.9 |

**Table 3.** The performance of supervised machine learning models (e.g., neural network, k nearest neighbour (kNN), and logistic regression) in comparison to 5 technicians. The k-fold cross validation sampling methods (e.g., k=2, 3, 5) and leave-one-out method were used to test and train the data. The performance of naïve Bayes model is well below the average human being (details in Supplementary Fig. 7). The abbreviations used were area under the curve (AUC), classification accuracy (CA), and F1-score is the harmonic mean for precision and sensitivity.

| Methods | AUC | CA | Sensitivity | Specificity | Precision | F1 |
| --- | --- | --- | --- | --- | --- | --- |
| neural network | 0.92 | 0.906 | 0.906 | 1 | 0.938 | 0.913 |
| kNN | 0.912 | 0.844 | 0.844 | 0.83 | 0.881 | 0.855 |
| logistic regression | 0.927 | 0.906 | 0.906 | 0.83 | 0.914 | 0.909 |
| average | 0.920 | 0.885 | 0.885 | 0.887 | 0.911 | 0.892 |
| Technician 1 | - | 0.781 | 0.778 | 0.800 | 0.955 | 0.857 |
| Technician 2 | - | 0.750 | 0.731 | 0.833 | 0.950 | 0.826 |
| Technician 3 | - | 0.813 | 0.846 | 0.667 | 0.917 | 0.880 |
| Technician 4 | - | 0.813 | 0.885 | 0.500 | 0.885 | 0.885 |
| Technician 5 | - | 0.813 | 0.815 | 0.800 | 0.957 | 0.880 |
| average | - | 0.794 | 0.811 | 0.720 | 0.932 | 0.866 |

The Hb genotyping identified single nucleotide polymorphism in the β-globin in the first and second samples, which was consistent with HbE (Fig. 5a) and HbD variant (Fig. 5b). A third rare Hb variant samples were identified with a G insertion at codon 27 of the β-globin gene (Fig. 5c). These haemoglobin variants exhibit similar clinical phenotype such as mild haemolysis and susceptible to oxidation^38,39^. The two-dimensional correlational mapping of Hb variants (Figs. 5a-c) revealed an unusual spectrum characteristic as compared to wild-type RBCs (Supplementary Fig. 3). The HbE variant (T_2_=1.06ms, T_1_=106ms), HbD variant (T_2_=1.20ms, T_1_=96.3ms), and the beta thalassemia variant (T_2_=1.40ms, T_1_=172ms) appears to have large and distorted T-peak with relatively short T_1_- and T_2_-relaxations as compared to wild-type Hb (T_2_=1.12ms, T_1_=188ms). The T-peak dispersion for the beta thalassemia variant with a mutated β-globin chain was particularly large with a flat plateau, suggesting that frame shift mutation causes a greater amount of haemoglobin instability^38^ (Fig. 5c).

In addition, the Hb variants appear to have much higher concentration of oxidized Hb as compared to the wild-type (Supplementary Fig. 3a). T_1_-relaxation stretching was observed for HbE variant (T_2_=0.94ms, T_1_=9.44ms) and the beta thalassemia variant (T_2_=0.56ms, T_1_=10ms), in agreement with commonly observed clinical phenotype such as mild haemolysis due to increased oxidative damage. Interaction of Hb variants and other forms of hemoglobinopathies can lead to complex thalassemia syndromes with varying clinical phenotypes (Fig. 2).

### Machine learning assisted medical decision

The 32 anonymized subjects consist of mixture of non-disease samples (wild-type), and disease samples (details in Supplementary Fig. 4b). The NMR correlational spectroscopy maps (molecular fingerprint) were converted into computer language for deep image analysis using statistical programming languages (e.g., *R*, Orange 3.1.2). Structural abnormalities in haemoglobin variants also lead to the observation of clinical methemoglobinemia in the late stage. The oxidized Hb samples were simulated examples for clinical methemoglobinemia.

The unsupervised learning techniques were used for dimension reduction (e.g., MDS), and classification (e.g., hierarchical clustering) to assist in making medical decision (Fig. 6). The 2D NMR correlational spectroscopy maps are complex 3D contour plots, and MDS technique was used to reduce higher dimension into two dimensional scatter plot which is more user-friendly for interpretation of information (Fig. 7a). Each feature (molecular fingerprint of one subject) was classified based on the common similarity within their intra-cluster as opposed to their inter clusters. Subjects were successfully classified into two clusters (disease (oxidized Hb, blue), non-disease (healthy wild type, red)) using the MDS technique (*P*-value<0.05), apart from the mutated counterpart (Hb variants, orange). In addition, the disease subtypes (sub-type 1: oxidized Hb, sub-type 2: partially oxidized Hb) were also observed (Fig. 7a). Distances between each subjects were shown in the heat map (Fig. 7b). Using hierarchical clustering, disease staging, prognosis or risk factor prediction (high/low risk factor) were enabled (Fig. 7c). Other techniques (e.g., Isomap, linearly local embedding, t-sne) were evaluated and similar results were reproduced qualitatively (Supplementary Fig. 5).

**Fig. 6:**
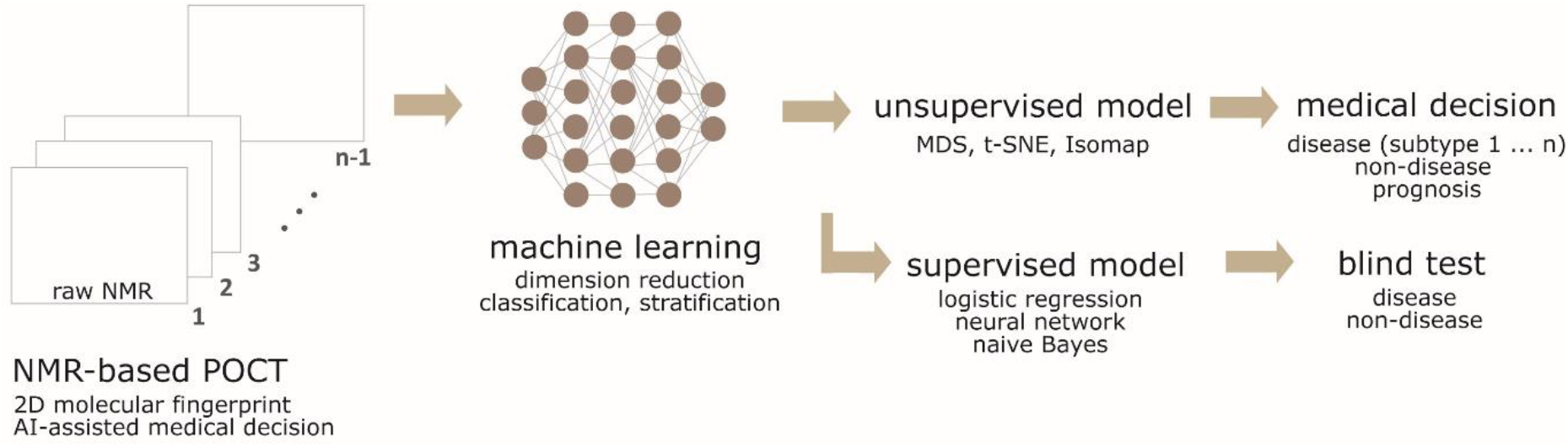
Machine-learning assisted NMR-POCT based medical decision. The workflow of machine learning in processing the complicated data into user-friendly medical decision (e.g., disease subtyping). The maps were converted into machine language using the image embedding (e.g., Squeeze Net) features. Dimensionality reductions were performed using various unsupervised models (e.g., MDS, t-SNE, Isomap). Supervised learning models (e.g., neural network, logistic regression, naïve Bayes) were used to train and predict the data. The performance of supervised learning techniques were compared to that of human performance (Table 3).

**Fig. 7:**
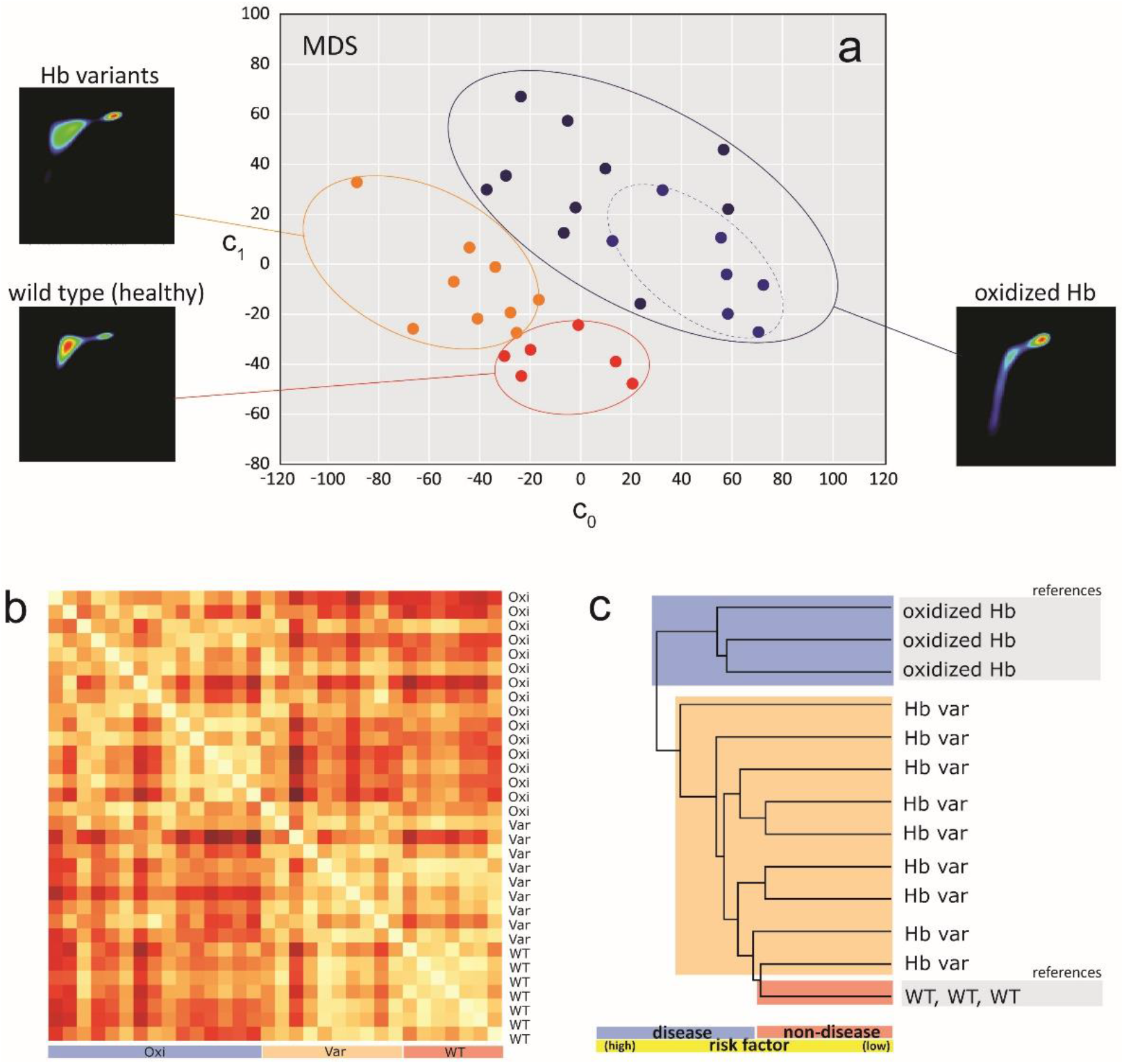
Machine-learning assisted NMR-based POCT in making medical decision. (a) The classification of three states (disease, non-disease, variants) and disease subtyping (sub-type 1: oxidized Hb, sub-type 2: partially oxidized Hb), and (b) heat map of 32 anonymized subjects processed using multidimensional scaling technique (300 max iterations, PCA-Torgersen). The legend (red, white) indicates (longer, shorter) distance between subjects. Other unsupervised models (e.g., linearly local embedding, Isomap, t-sne) were also evaluated for comparison (Supplementary Fig. 5). (c) The hierarchical clustering enabled disease staging, prognosis or risk factor prediction (high/low risk subject) with respect to standard reference. For simplicity, three referencing states (WT and oxidized Hb) were shown. The non-disease state consists of (healthy wild-type), and disease state consist of (oxidized Hb, Hb variants). The short forms used were wild type (WT), oxidized Hb (Oxi), and Hb variants (Var). The clustering circles (dotted lines) were drawn for eye-balling purposes. The NMR correlational maps of each subjects is shown in Supplementary Fig. 4.

### Blinded test: machine vs human learning

The 32 anonymized subjects consist of mixture of non-disease samples (wild-type) and disease samples (details in Supplementary Fig. 4b). Supervised learning models (e.g., logistic regression, neural network, k nearest neighbours (kNN) and naïve Bayes) were used to evaluate its’ efficiency against human-being. K-fold cross validation (e.g., k=2, 3, 5) and leave-one-out method were used for samplings. Five technicians were trained to differentiate between (diseases, non-disease) and subsequently were asked to classify the state of the spectrum based on a binary decision (diseases, non-disease) in blinded manner. At the end of the experiment, the results were cross-checked and classified as true positive, true negative, false positive and false negative (Supplementary Fig. 6). On-average, the machine learning models (e.g., CA=0.885, sensitivity=0.885, specificity=0.887) outperformed the human being (e.g., CA=0.794, sensitivity=0.811, specificity=0.720) in many aspects, when k=5 (Table 3). The performance of the supervised models, in general, improved with increasing value of k and achieved the maximum point when ‘leave-one-out’ method was used in training the datasets (details in Supplementary Figure 7). Noticeably, the performance variation between each individual was larger than that of machine learning models as a result of human subjective judgment. On-average, machine learning models (30 seconds) also took much shorter time than human (about 10 minutes) to complete the tasks given.

## Discussion

In this work, we showed that detailed and specific molecular microenvironment of water-proton interactions in blood can be mapped out using the two-dimensional T_1_-T_2_ correlational spectroscopy. Interestingly, as water is ubiquitous to life form, water-protein interactions (e.g., the protein hydration) attracted considerable interests from terahertz spectroscopy^40^ to neutron scattering^41^, provides an equivalent of ‘inverse proteomic’ information. This adds a new dimension to the existing traditional omics framework (e.g., genomic, proteomic) potentially revealing many biological pathways and understanding of fundamental of biological processes which have never been examined before.

It is demonstrated that the proposed technique here is capable of rapid label-free phenotyping the biological fluids in various physiological conditions (e.g., de/oxygenation level) and pathological states (e.g., blood oxidation, hemoglobinopathies) in uniquely personalized manner. We showed that time-to-result could be accomplished in minutes (Supplementary Fig. 1). With the recent availability of ultrafast signal acquisition methods^11^ and efficient inversion algorithm^7^, real-time characterization and monitoring is possible. Aided with machine learning techniques, complicated NMR correlational maps were immediately transformed into clinically meaningful and user-friendly information.

Secondly, encoding multidimensional biochemical and biophysical information at molecular level using two-dimensional relaxation profiling (instead of chemical shifts), circumvent the limitation of using conventional big footprint NMR. Unlike high field NMR spectroscopy, mass spectrometry, high performance liquid chromatography where the instrumentation are often bulky and expensive (Table 4), an interesting NMR-based point-of-care technology (POCT) alternative proposed in this work offers inexpensive assay and instrumentation (e.g., open source code software-defined-radio^42–44^). Importantly, the unique and specific molecular fingerprint of liquid biopsy is able to provide a multiple global snapshot for disease dynamic monitoring in a minimally invasive manner^45,46^.

**Table 4.**
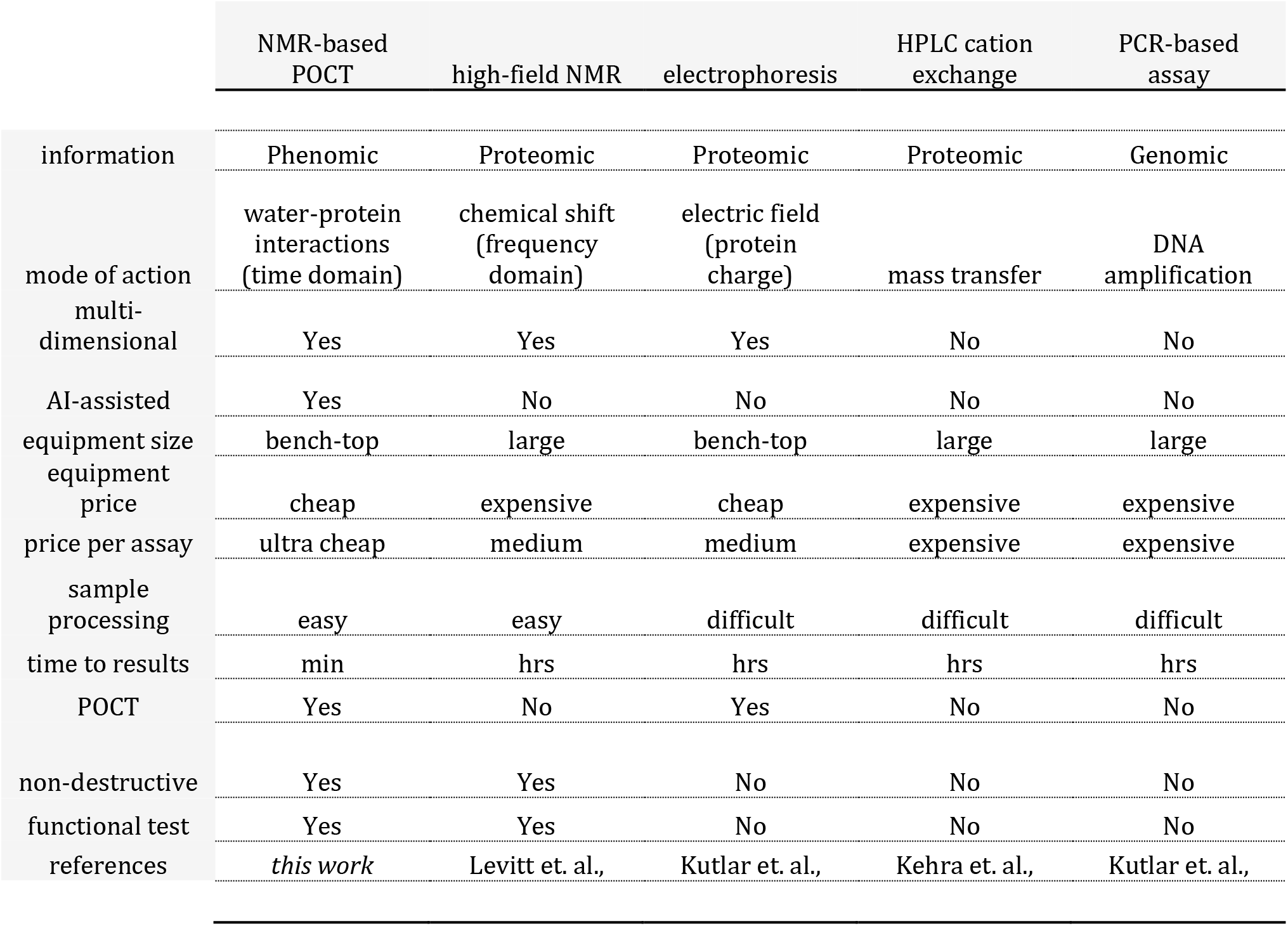
Comparison between the current proposed NMR-based POCT with AI-aided technology and existing competing technologies (e.g., high-field NNR^48^, electrophoresis^49^, hplc^50^, and PCR-based assay^45^) which have been reported for clinical hemoglobinopathies. The price per assay for NMR-based POCT refers to a single microcapillary tube (<$0.10). Furthermore, this method is label-free and therefore no chemical treatment is required. The abbreviation for HPLC: high performance liquid chromatography.

In summary, a novel concept of NMR-based POCT rapid and label-free molecular phenotyping of biological fluids were introduced. The use of machine learning algorithm improves the delivery of information (e.g., speed and accuracy), which may become a key factor in speeding up the translational technological innovations to clinical routine and practices. Assessment of multidimensional relaxation components of the blood was shown to be highly time- and patient-specific, delivering personalized information that is critical in clinical diagnostic, monitoring and prognostic purposes.

## Author Contributions

W.K. Peng conceived the original idea, wrote the first draft of the paper, designed the experiments/protocols (e.g., NMR correlational experiments, machine learning coding), built the entire hardware setup and performed micro MR measurements. T-T Ng and Lim Daniel contribute in modifications on MATLAB code for ILT analysis. T.P. Loh kindly provide the blood samples and engaged in various discussion on translational clinical aspects.

## Acknowledgement

This research was supported by the International Iberian Nanotechnology Laboratory (INL Start Up Grant and INL Seed Grant). T-T Ng acknowledges the support of A*STAR AGA that provides access to a pool of very talented interns through its student attachment programme. W.K. Peng would like to personally thank L. Daniel, an internship student along with T-T. Ng ’s co-supervision. L. Daniel conducted various preliminary MR measurements, and modifying the ILT code using MATLAB. We acknowledge Chin Hin Ng of Department of Oncology, National University of Hospital, Singapore for assistance in obtaining approval from the National Health Group (NHG) Institution Review Board.

## Methods Online

### NMR setup and parameters

The ^1^H magnetic resonance measurements of packed bulk red blood cells were carried out at the resonance frequency of 21.57 MHz using a portable permanent magnet (Metrolab Instruments, Switzerland), B_o_=0.5T using a benchtop-type console (Kea Magritek, New Zealand). A temperature controller was set to maintain the measurement chamber at 24.5°C. The T_1_-T_2_ correlational pulse sequences were set at standard inversion recovery, followed by Carr-Purcell-Meiboom-Gill (CPMG) train pulses (Fig. 1).

The experimental parameters used; echo time=200 μs, number of echoes=2000 (for oxidized, and deoxygenated state) and 4000 (for oxygenated state), T_1_ incremental steps=32 (logarithmic) steps, and signal averaging=4. A recycle delay of 2s was set between each experiment to provide sufficiently long time to allow all the molecular spins to return to thermal equilibrium. The total acquisition time depends on the combination of a number of factors (e.g., number of scans, T_1_-incremental).

We demonstrated that a total experimental time in less than 6 minutes is sufficient for a high sensitivity and good spectral resolution, and without losing the spectral integrity (details in Supplementary Fig. 1). The 2D correlation maps were processed using built-in ILT algorithm (FISTA inversion)^47^ method with 5000 iterations and smoothing parameter of 1 were used. The inversion typically completed in less than 2 minutes using a desktop computer (Intel Core Pentium i3 CPU @ 3.2GHz, 1.74Gb RAM).

### Clinical ethics and protocols

This study received ethics approval from the local Institutional Review Board of the National Healthcare Group. K2 EDTA-anticoagulated whole blood samples were washed and re-suspended with phosphate buffer saline (PBS). All blood samples were either used immediately or kept at 4°C and used within three to four days (unless mentioned otherwise) of collection before the micro MR analysis. To induce the Hb into various derivative states, the blood samples were incubated with the desired chemical as mentioned in the *Text* (e.g., sodium nitrite) and finally washed to remove the chemical residual. Heparinized micro capillary tubes (Fisher Scientific, PA) were used to transfer the processed blood and finally spun down at 6000*g* for 1 minute to obtain packed red blood cells for MR measurements.

### Machine learning algorithm and workflow

The NMR-based POCT can be used with or without the assistant of AI (Fig. 2). Machine learning techniques were used to transform the human complicated data (e.g., 2D NMR correlational maps) into user-friendly medical decision making following the workflow developed (Fig. 6). The maps were converted into machine language using the image embedding features (e.g., Squeeze Net). Machine learning techniques were used to perform dimension reduction using various techniques (e.g., MDS, t-SNE, Isomap) (Supplementary Fig. 5).

### Blinded test

Supervised learning models (e.g., neural network, k nearest neighbor, logistic regression, and naïve Bayes) were used to train and predict the data. We first trained 5 human beings to differentiate between (diseases, non-disease) and asked them to classify 32 anonymized subjects that were not seen before (Supplementary Fig. 4). They were allowed to backtrack (and change) the results as long as it was within the allocated time-frame (10 minutes). At the end of the experiment, the results were cross-checked and classified them as true positive (TP), true negative (TN), false positive (FP) and false negative (FN) (Supplementary Fig. 6). Statistical programming languages (e.g., Orange 3.1.2) was used for machine learning algorithm running on a personal laptop (Intel Core Pentium i7 CPU @ 2.70GHz, 8.00 GB RAM). Once the models in machine learning were built, the run test takes less than 30 seconds to complete all the tasks, while each of the human beings took about 10 minutes on-average.

### Statistical methods

Two tailed Student’s T-test was used to calculate the *P*-value.

## Data availability statement

The machine learning algorithms and 2D NMR raw maps are available upon reasonably request at.

